# SPARC mediates tumour–stroma intercellular communication through endosomal regulation of Delta and Notch signalling

**DOI:** 10.64898/2026.07.04.736483

**Authors:** Aurélien Guillou, Nourhene Ammar, Olivier Josse, Emma Leroux, Tsveta Kamenova, Torcato Martins, Samuel Delage, Maurice Joseph Ringuette, Sarah Bray, Hadi Boukhatmi

**Author notes:** Equal contribution (Nourhene Ammar, Olivier Josse).

## Abstract

Tumour progression relies on reciprocal communication between genetically altered cancer cells and surrounding stromal cells. While the genetic alterations that initiate tumorigenesis have been extensively studied, the dysregulated feedback signalling provided by co-opted stromal cells remains poorly understood. Here, we used a *Drosophila* cancer model to address this question and identified the matricellular protein SPARC as a mediator of tumour–stroma communication. SPARC is produced by mesenchymal cells and transferred into epithelial tumour cells, where it is internalized through the endocytic pathway. Following uptake, SPARC accumulates in Rab7 positive late endosomes and colocalize with the Notch ligand Delta. SPARC internalization promotes endosomal enlargement and reduces endosome dynamics. Increased SPARC levels in epithelial tumours indirectly attenuate Notch signalling activity through at least altered Delta trafficking. We further identify the N-terminal acidic domain of SPARC as specifically required for its targeting to Delta-associated endosomes. Together, our findings uncover a stromal feedback mechanism by which SPARC modulates Notch signalling through endosomal regulation during tumour development.

## Introduction

Cells continuously communicate with their neighbours, and the signals exchanged are crucial for the organization and maintenance of multicellular tissues. Tight regulation of these signalling processes ensures coordinated growth, patterning, and tissue homeostasis. In contrast, aberrant activation or inappropriate deployment of these pathways underlies many diseases, including cancer [1]. In ovarian carcinomas, for example, reciprocal signalling between genetically altered epithelial cells and surrounding stromal cells critically supports tumour progression and malignancy [2–4]. Untransformed mesenchymal stromal cells are thought to be “educated” by the cancerous cells, acquiring tumour-supportive properties that contribute to tumour development [5]. While the genetic mutations that initiate tumorigenesis have been extensively characterized, an important challenge is to identify the signals produced by stromal cells and to elucidate how they sustain tumour growth and reinforce malignant behaviour. Understanding these mechanisms is crucial and may reveal new opportunities for alternative therapeutic strategies.

We investigate the role of the Secreted Protein Acidic Rich in Cysteine (SPARC) in tumor–host communication. SPARC is a secreted matricellular glycoprotein that modulates cell–cell and cell–matrix interactions and is dynamically regulated during development, tissue remodelling, and injury [6]. As a member of the matricellular protein family, SPARC does not primarily provide structural support to the extracellular matrix but instead influences cellular behaviour by regulating adhesion, proliferation, and responses to extracellular cues [7]. SPARC is also implicated in tumour development, yet its role in cancer remains complex and context dependent. It has been shown that SPARC can either restrain or promote tumour growth and metastasis, depending on tissue type and tumour context [8, 9]. These dual effects suggest that SPARC does not act as a simple oncogene or tumour suppressor but instead functions as a context-dependent regulator of tumour–microenvironment interactions. Despite extensive investigation, the mechanisms by which SPARC exerts these diverse effects remain limited. In particular, how SPARC produced by genetically normal stromal or mesenchymal cells influences tumour cells behaviour, and how such signals are transmitted at the cellular level, is still unclear.

*Drosophila* genetic models have been instrumental in dissecting molecular mechanisms underlying intercellular communication in cancer [10, 11]. One such model is the *EGFR-psq^RNAi^*tumour model, in which the epidermal growth factor receptor (EGFR) signaling is overactivated in the wing disc epithelium using the Apterous driver (Ap-Gal4) in combination with RNAi-mediated depletion of the chromatin regulator *pipsqueak* (*psq*) ([12] and Figure S1A). In this model, tumours grow as a large multilayered mass composed of transformed epithelial cells intermingled with genetically normal myoblasts that form the mesenchymal compartment (Figure S1A-B’). This model provides a simple and genetically tractable system to study tumour–stroma interactions *in vivo*. Indeed, the transformed epithelial cells induce proliferation and maintenance of adjacent myoblasts through, at least, Notch signalling, resulting in coordinated expansion of both the epithelial and mesenchymal compartments to comparable sizes [12, 13]. The Notch receptor is present in both epithelial and mesenchymal cells (Figure S1A–B’), whereas the Notch ligand Delta is strongly expressed in epithelial tumour cells (Figure S2C–F’). Epithelial Delta activates Notch signalling in neighbouring mesenchymal cells through direct cell–cell interactions and cytoneme-mediated contacts. This signalling promotes expansion of the tumour-associated mesenchyme and coordinates its growth with the epithelial compartment [13]. The mesenchymal cells are in turn required to sustain epithelial growth and tumour progression through a paracrine signalling loop [12, 14]. However, the identity of the mesenchyme-derived signals that sustain epithelial tumour growth, and the mechanisms by which they act on tumour cells, remain poorly understood.

Here, using the *EGFR-psq^RNAi^* model, we show that SPARC mediates tumour–stroma communication by regulating endosomal trafficking. SPARC is produced by mesenchymal cells and transferred into epithelial tumour cells through the endocytic pathway. Once internalized, SPARC accumulates in Rab7-positive late endosomes, where it colocalizes with the Notch ligand Delta. Importantly, SPARC accumulation within these compartments is associated with endosome enlargement and reduced motility. These alterations in endosomal trafficking indirectly attenuate Notch signalling in epithelial tumours, likely by shifting Delta-containing endosomes toward late endosomal compartments at the expense of recycling to the plasma membrane. Finally, we identify the N-terminal acidic domain of SPARC as specifically required for its targeting to endosomes and for its effects on endosome dynamics. Together, these findings reveal an intracellular mechanism by which SPARC modulates endosomal control of Notch signalling during tumour development.

## Results

### SPARC is expressed in the tumour associated mesenchyme

To investigate whether SPARC may contribute to signalling between epithelial tumours and mesenchymal cells, we first analysed its expression in wild-type wing discs. SPARC expression was detected in wing disc–associated myoblasts, as identified by the myoblast markers Cut and CG9650 (Figure 1A–C, Figure S2A–B″ and Video S1). These cells correspond to adult muscle progenitors (AMPs), which are associated with the wing disc epithelium during larval stages and ultimately give rise to the adult flight muscles [15, 16]. This pattern was independently confirmed using two reporter strategies: (i) the SPARC-Gal4 (SPARC^[MI00329-GAL4]^) driver to express a UAS membrane marker (UAS-mCD8–RFP), and (ii) the BM40-SPARC-GFP fosmid reporter line [17]. Both reporters label wing disc myoblasts (Figure 1D–E’’). Importantly, the BM40-SPARC-GFP cassette is functional, as it rescues the lethality caused by *sparc* loss-of-function allele (Figure S3 and Materials and Methods). By contrast, SPARC is not expressed by the epithelial wing disc cells, as neither SPARC-Gal4 activity nor *sparc* mRNA was detected in this compartment (Figure 1D-F and Figure 2 C-C’ and Video S1). We next analysed SPARC expression in *EGFR-psq^RNAi^* tumours and found that it is maintained in the mesenchymal cells associated with the tumours (Figure 1G–H’’). Quantification of *sparc* transcripts revealed a significant upregulation in tumours compared to wild-type wing discs, suggesting a potential role in tumour development (Figure 1I). Altogether, these data identify SPARC as a mesenchyme derived factor upregulated during tumorigenesis, prompting us to investigate its functional contribution to tumour-stroma communication.

**Figure 1.**
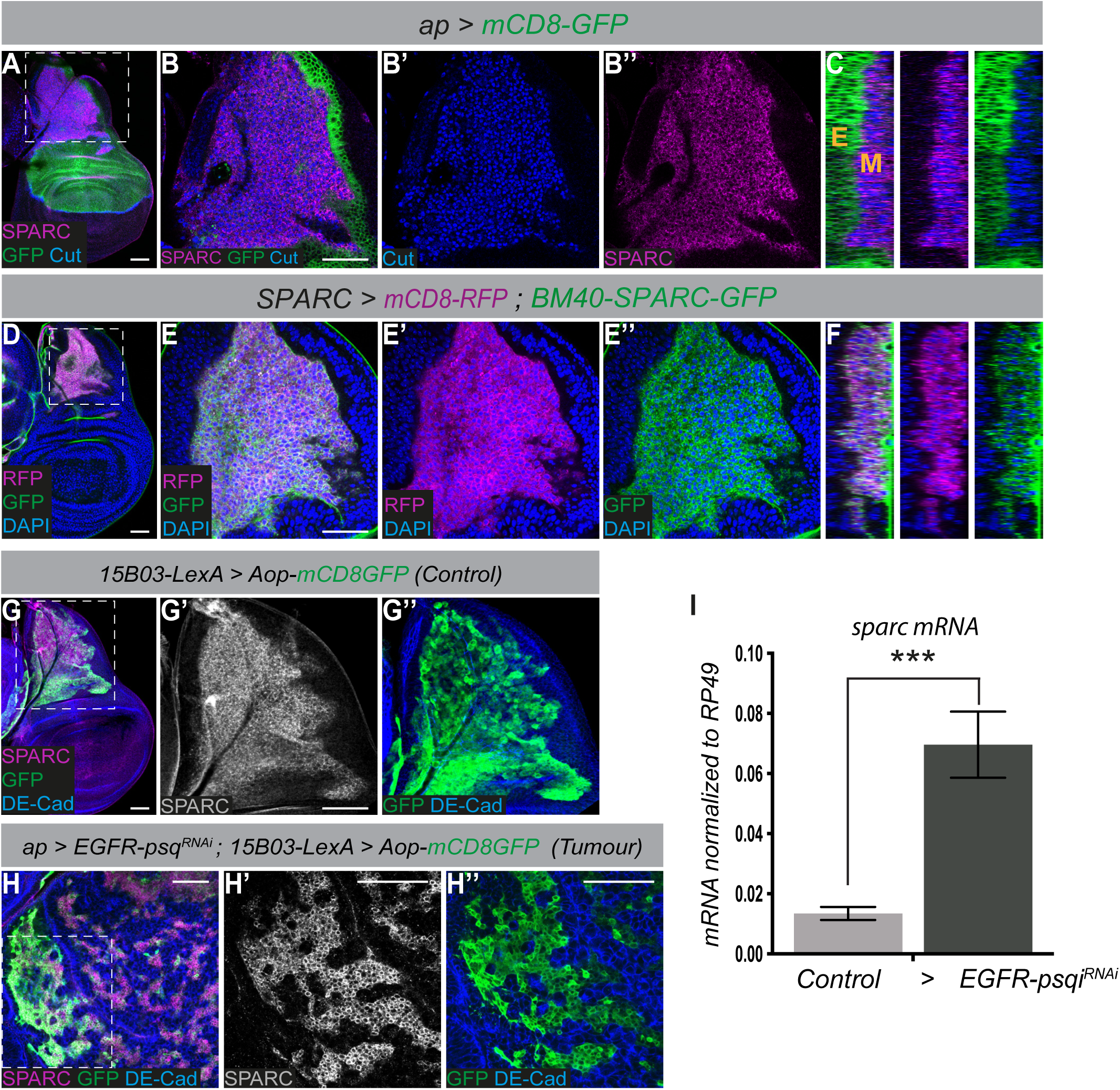
SPARC is detected in tumour mesenchymal cells. **(A-B).** *Apterous-Gal4* (green, *ap-Gal4, UAS-mCD8-GFP*) drives expression in epithelial cells but not in the myoblasts (Cut, blue). **(B’-B’’).** SPARC (magenta) is expressed in myoblasts (Cut, blue). **(C).** Orthogonal views of (B) showing that SPARC expression (magenta) is restricted to myoblasts (M) and absent from the epithelium (E). **(D-E’’).** *SPARC-Gal4* (magenta, *SPARC > mCD8–RFP*) drives expression in myoblasts that express *BM40-SPARC-GFP* (green). DAPI (blue) marks all nuclei. **(F).** Orthogonal views of (E) showing both *SPARC > mCD8–RFP* (magenta) and BM40-SPARC-GFP expression (green) in the myoblasts (M). **(G-H’’).** Expression of SPARC (magenta in G and Gray in G’) is detected in myoblasts (G-G’’) and in the mesenchymal cells of the *EGFR-psq^RNAi^* tumours (H-H’’). 15B03-LexA (green, *15B03-LexA; LexAop-mCD8-GFP*) drives expression in mesenchymal cells. DE-Cadherin (DE-Cad; blue) marks epithelial cells. **(I).** *sparc* mRNA expression levels measured by quantitative RT-PCR. RNAs were prepared from three independent experiments. Error bars represent s.e.m. Scale bars: 50μm.

**Figure 2.**
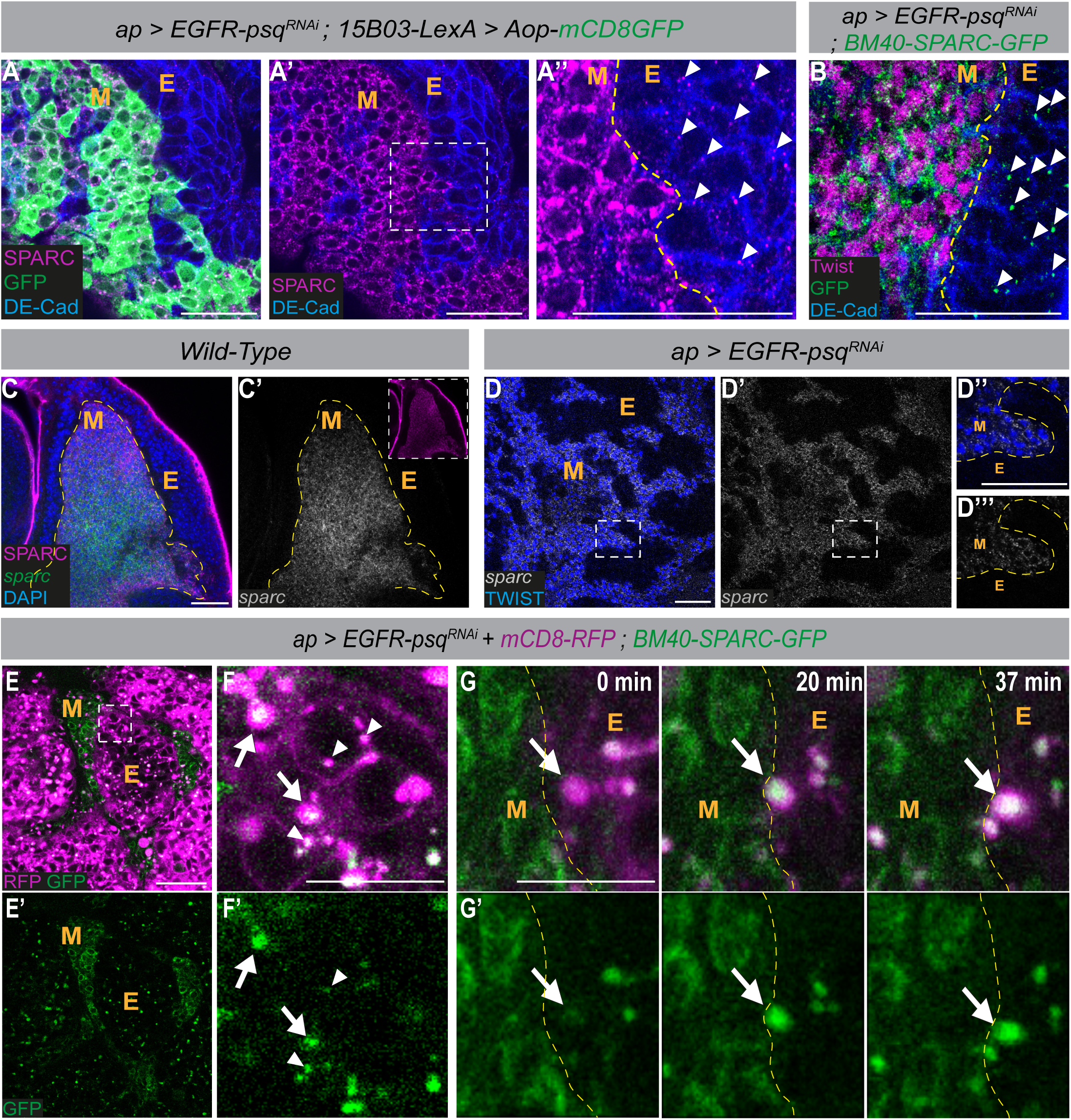
Mesenchymal SPARC is transferred to tumour epithelial cells. **(A-A’’).** Expression of SPARC (magenta) is detected in both the mesenchymal cells (green, *15B03-LexA; LexAop-mCD8-GFP*) and in epithelial cells visualized by DE-Cad (blue). **(B).** Expression of BM40-SPARC-GFP line (green) is detected in both mesenchymal cells (M) (magenta, Twist) and in epithelial cells of the *EGFR-psq^RNAi^*tumours (E) marked by DE-Cad (blue). Yellow dashed lines delimit Mesenchymal cells (M) and Epithelial cells (E). Scale bars: 25μm. **(C-D’).** Detection of *sparc* transcripts (gray) by smFISH in wild-type myoblasts (C-C’) and mesenchymal tumours (D-D’’’). SPARC (magenta, C) and Twist (blue, D) label the wild-type myoblasts and the mesenchymal cells, respectively. Scale bars: 25μm. **(E-E’).** Single time point from a time-lapse sequence (see Video S2) of *EGFR-psq^RNAi^* tumour sample showing BM40-SPARC-GFP (green) and epithelial cells (magenta, *ap > mCD8-RFP*). Scale bar: 25μm. **(F-F’)** higher magnification of the boxed region in (E). Large and small SPARC containing vesicles are indicated by arrow and arrowhead, respectively. Scale bar: 10 μm. **(F).** Time course (min) of the *EGFR-psq^RNAi^* tumours showing the internalization of SPARC (green, BM40-SPARC-GFP, arrows) from the mesenchymal compartment to the epithelial vesicles. Scale bar: 10μm.

### SPARC is produced by mesenchymal cells and internalized by epithelial tumour cells

Given that SPARC is a secreted protein, we asked whether it could act as a signalling molecule that is involved in the signalling between mesenchymal and epithelial cells. Higher-magnification analysis of *EGFR-psq^RNAi^* tumours shows that, in addition to mesenchymal SPARC expression, SPARC can also be detected in epithelial tumours, as marked by DE-Cad (Figure 2A–A’’). In mesenchymal cells, SPARC is robustly detected throughout the cell, whereas in epithelial tumour cells it instead localizes to discrete, round intracellular and membrane associated puncta (Figure 2A’’). This pattern was independently confirmed using the BM40-SPARC-GFP line (Figure 2B and Figure S2C-C’’). To analyse whether the SPARC detected in epithelial tumours originates from mesenchymal cells rather than being transcribed by the epithelial tumour cells themselves, we examined *sparc* transcripts expression using single-molecule FISH (smFISH, [18]) in both wild-type discs and *EGFR-psq^RNAi^*tumours (Figure 2 C-D). We found that *sparc* is exclusively transcribed in the mesenchymal cell population in both wild-type and *EGFR-psq^RNAi^*tumours, with no detectable transcription in epithelial cells (Figure 2 D’-D’’’ and Figure S2D-E). These observations suggest that the SPARC detected in epithelial tumour cells may originates from neighbouring mesenchymal cells. To test this hypothesis, we performed extended live imaging of *EGFR-psq^RNAi^* tumours in which epithelial cells were labelled with membrane-targeted RFP and SPARC was visualized using the BM40-SPARC-GFP line (Figure 2E–F′ and Video S2). We found that, in epithelial tumours, SPARC positive puncta corresponded to highly mobile intracellular vesicles, displaying variable sizes and motility characteristics. Higher-magnification analysis of these time lapses revealed that SPARC is taken up from the mesenchymal compartment into epithelial cells through these vesicular structures at the interface between these two compartments (Figure 2G-G’ and Video S3). These data indicate that SPARC present in the epithelial tumour compartment does not result from local production but is instead incorporated from adjacent mesenchymal cells via intracellular vesicles.

### SPARC exerts distinct pro- or anti-tumor functions depending on its cellular localization

Given that SPARC is detected in both mesenchymal cells and epithelial tumour cells, we next aimed to define its functional role in each cellular compartment. To specifically assess the requirement for mesenchymal SPARC, we used the LexA system [19] to selectively downregulate *sparc* in mesenchymal cells via 15B03-LexA driver. Targeted depletion of *sparc* in the mesenchyme resulted in a strong inhibition of tumour growth (Figure 3 A-C), indicating that SPARC is required in this compartment to support tumour expansion. To determine the functional consequence of SPARC following its transfer into epithelial tumour cells, we overexpressed SPARC specifically in the epithelial compartment and measured tumour size. This approach allowed us to assess the cell-autonomous effects of epithelial SPARC without altering SPARC levels in the mesenchymal compartment. Epithelial SPARC overexpression led to a significant reduction in tumour growth compared with control tumours (Figure 3D–F). Together, these results show that SPARC has distinct effects on tumour growth depending on its cellular localization. In mesenchymal cells, SPARC promotes tumour growth, whereas in epithelial tumour cells SPARC acts to restrain tumour expansion. We focused subsequent work on the role of SPARC within epithelial tumour cells, aiming to elucidate how mesenchyme-derived SPARC functions as an intercellular signal to modulate epithelial tumour behaviour.

**Figure 3.**
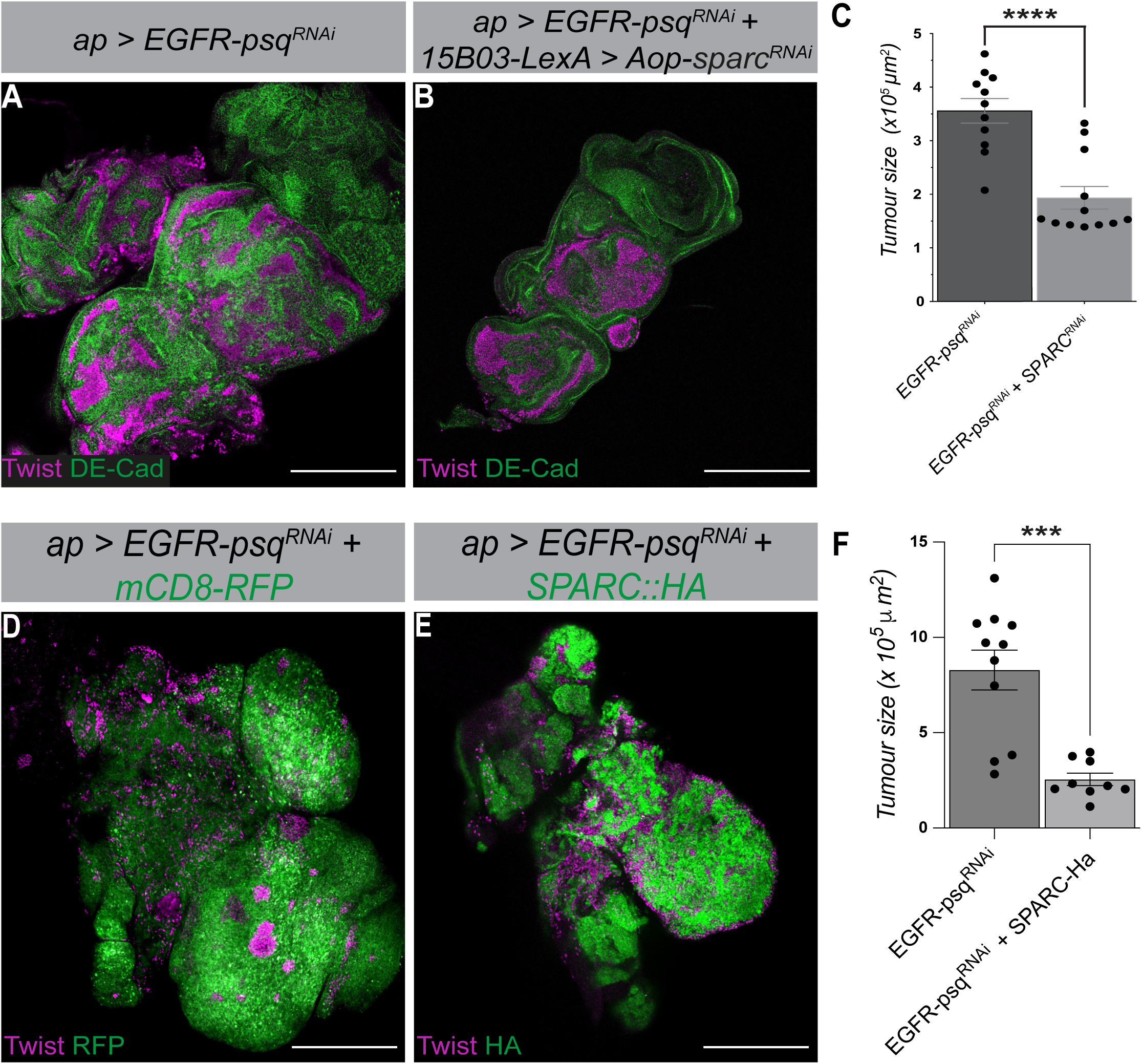
SPARC exerts distinct functions in the mesenchymal and epithelial compartments. **(A-B).** Specific depletion of *sparc* in the mesenchymal cells reduces tumour growth. *ap > EGFR-psq^RNAi^* tumours in control conditions (A) and after mesenchymal knockdown of *sparc* using *15B03-LexA; LexAop-sparc^RNAi^*(B). DE-Cadherin (green) marks epithelial tumor cells and Twist (magenta) labels mesenchymal cells. **(C).** Quantification of tumour size for the genotypes shown in (A) and (B). *\*\*\*\*p* < 0.0001 (unpaired t test; n = 11 and 12 tumours for control and *sparc^RNAi^*, respectively; data pooled from three independent experiments). **(D-E).** Epithelial overexpression of SPARC restricts tumor growth. Confocal images of *ap > EGFR-psq^RNAi^* tumours in control conditions (D) and upon epithelial expression of UAS-SPARC::HA (E). **(D).** RFP (green) indicate *ap-Gal4* expression in the epithelial cells (D). **(E).** HA (green) indicates ectopic expression of SPARC::HA. Twist (magenta) marks mesenchymal cells. **(F).** Quantification of tumour size for the genotypes shown in (D) and (E). *\*\*\*p* < 0.001 (unpaired t test; n = 11 and 9 tumours for control and SPARC::HA, respectively; data pooled from two independent experiments). Scale bars: 200μm.

### Ectopic SPARC attenuates Notch signalling in epithelial tumours

During tumour development, multiple developmental signalling pathways are co-opted to regulate growth [12, 13]. We asked whether mesenchymal SPARC interferes with such pathways in the epithelial compartment. As a first step to test this hypothesis, we overexpressed SPARC in the wing disc epithelium using *Ap-Gal4* and assessed its impact on adult wing development. SPARC overexpression induced a wing phenotype that phenocopied Notch mutants in approximately 10% of the hatched adult flies (Figure 4A–D). These data indicate that SPARC may interfere with Notch pathway activity in epithelial cells. The dorsoventral (D/V) boundary of the wing pouch is a classical system for studying Notch activity [20]. To monitor variations in Notch signaling in tumour epithelial cells, we used the NRE-RFP (Notch Responsive Element) reporter, which is normally expressed at the D/V boundary and responds sensitively to alterations in Notch pathway activity (Figure 4E-E’’ and [20]). We focused our analysis on 4-day-old tumours, when the pouch compartment remains morphologically distinguishable from other tumorigenic cells. In wild-type discs, NRE-RFP expression is restricted to the D/V boundary of the pouch (Figure 4E–E″). In contrast, *EGFR-psq^RNAi^* tumours show strong upregulation of NRE-RFP throughout the epithelial cells of the pouch, indicating elevated Notch signalling activity (Figure 4F–F″ and H). This observation is consistent with the high levels of Delta detected in epithelial tumour cells (Figure S1) and with our previous finding that epithelial Delta is required for tumour growth [13]. Strikingly, overexpression of SPARC in epithelial tumour cells significantly reduced NRE-RFP levels compared to *EGFR-psq^RNAi^* tumours alone, although Notch activity was not fully restored to control levels (Figure 4G–H). Consistent with these observations, NRE-RFP levels are higher in tumour regions depleted of mesenchymal cells, whereas epithelial regions enriched in mesenchymal cells display reduced reporter activity (Figure 4I–K). Together, this suggests that SPARC originating from the mesenchymal compartment may locally dampen Notch activity in epithelial tumour cells.

**Figure 4.**
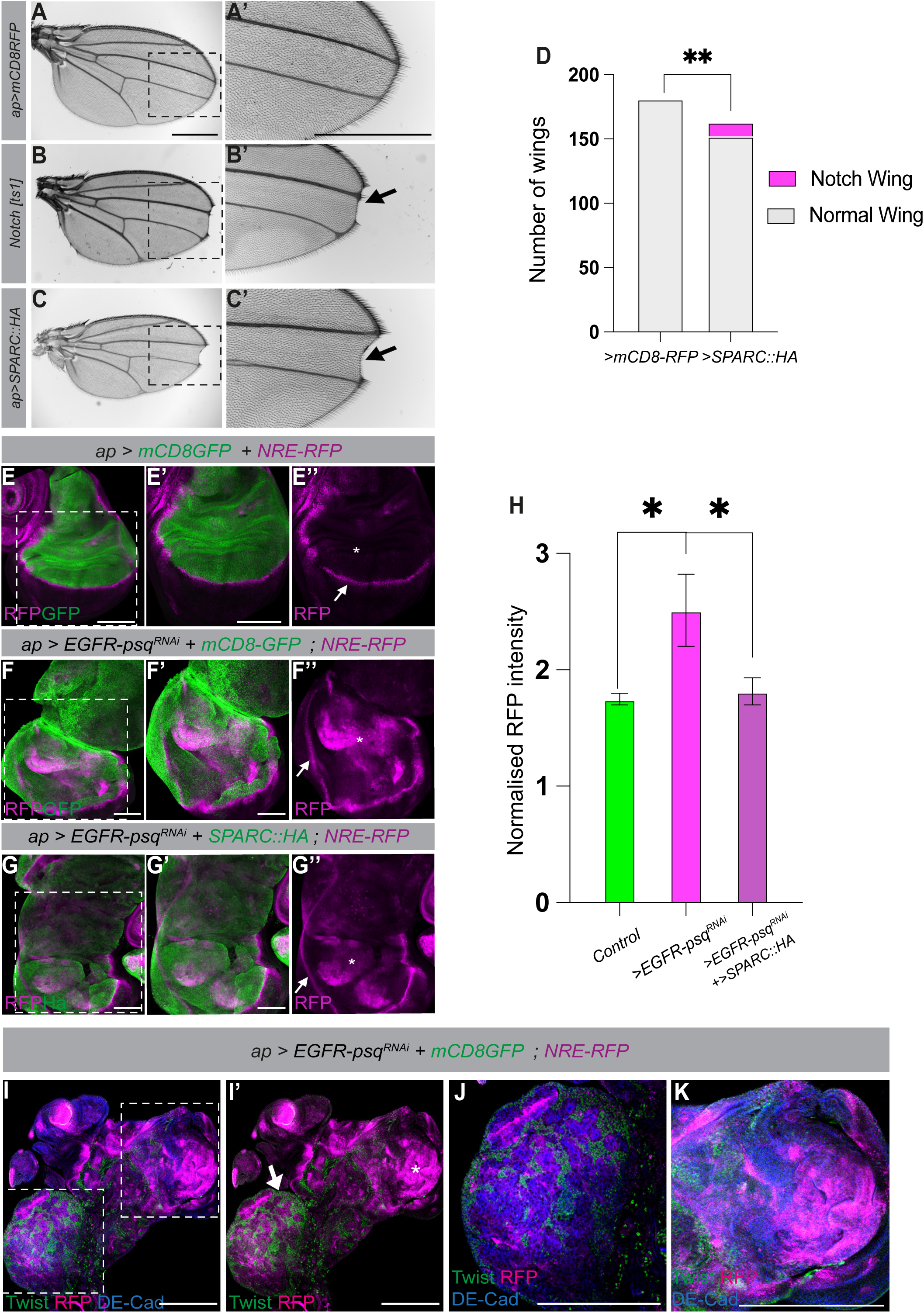
SPARC attenuates Notch signaling activity in epithelial tumours. **(A-C).** Adult wings from control flies (*Aap>mCD8-RFP*, A–A′), a Notch loss-of-function (*Notch [ts1],* B–B′), and flies overexpressing SPARC in the wing disc (*ap > SPARC::HA*, C–C′). Arrows indicate wing margin defects characteristic of reduced Notch signalling. Dashed boxes correspond to magnified views in A′–C′. Scale bars: 500μm. **(D).** Quantification of wing phenotypes showing an increased frequency of Notch margin defects upon SPARC overexpression. *\*\*p* < 0.01 (Chi-square test; n = 181 and 152 wings for control and SPARC::HA, respectively; data pooled from four independent experiments). **(E-E″).** Notch pathway activity in control wing discs visualized using the Notch responsive reporter NRE-RFP (magenta). The dorsal compartment of the disc is marked by mCD8-GFP (*ap>mCD8-RFP*, green). NRE-RFP expression is restricted to the dorsoventral (D/V) boundary (arrow). **(F-F″)** In *EGFR-psq^RNAi^*tumours, NRE-RFP expression is upregulated throughout the pouch region (asterisk). **(G-G″).** Overexpression of SPARC in epithelial tumour cells (*EGFR-psq^RNAi^*+ *SPARC::HA*) reduces NRE-RFP levels compared to tumours alone. **(H).** Quantification of normalized NRE-RFP fluorescence intensity in the indicated genotypes. *\*p* < 0.1 (unpaired t test, n=13 control wing discs, n=10 *EGFR-psq^RNAi^*wing disc, n=12 *EGFR-psq^RNAi^* + *SPARC::HA* wing discs; data pooled from four independent experiments). Scale bars: 100μm. **(I-I’).** NRE-RFP (magenta) expression in the *EGFR-psq^RNAi^* tumours. Twist (green) marks mesenchymal cells, DE-Cad (bleu) marks the epithelial cells. (I’) NRE-RFP levels are higher in tumour regions lacking mesenchymal cells. The asterisk indicates regions with high NRE-RFP signal, whereas arrows indicate mesenchyme-rich regions where NRE-RFP levels are reduced. **(J-K).** higher magnifications of the boxed regions in (I). Scale bars: 200μm.

### Transferred SPARC accumulates in Rab7 endosomes containing Delta

To understand how SPARC reduces Notch activity in epithelial tumour cells, we sought to identify the cellular compartment where SPARC and components of the Notch pathway may converge. As an initial step, we overexpressed SPARC in wild-type wing discs using the Ap-Gal4 driver and analysed its distribution (Figure 5A–F). Ap-Gal4 is activated in a restricted epithelial subdomain of the wing disc that abuts the dorsoventral (D/V) boundary, which is characterized by Delta expression (Figure 5A–B’). This strategy allows SPARC to be expressed in a restricted epithelial domain while enabling analysis of its localization in neighbouring Delta expressing cells. We found that transferred SPARC localized to Delta positive puncta (Figure 5D–F’’). These puncta predominantly correspond to late endosomes, as indicated by colocalization with the late endosomal marker Rab7 (Figure 5C–C’’). Consistently, the analysis of SPARC in *EGFR-psq^RNAi^* tumours revealed a similar pattern (Figure 5G–G’’’). Indeed, SPARC is predominantly detected in Rab7 endosomes within the epithelial compartment (89%, 128/144 vesicles analysed) (Figure 5G–G’’’). Importantly, Delta is also enriched within these Rab7/SPARC late endosomes (Figure 5H–H‴). By contrast, analysis using the recycling endosomal marker Rab11 revealed no detectable association between SPARC and Delta in Rab11 positive recycling endosomes (Figure 5I–I’’’). Together, these data indicate that following uptake into epithelial cells, SPARC is routed to Rab7 compartments that also contain Delta.

**Figure 5.**
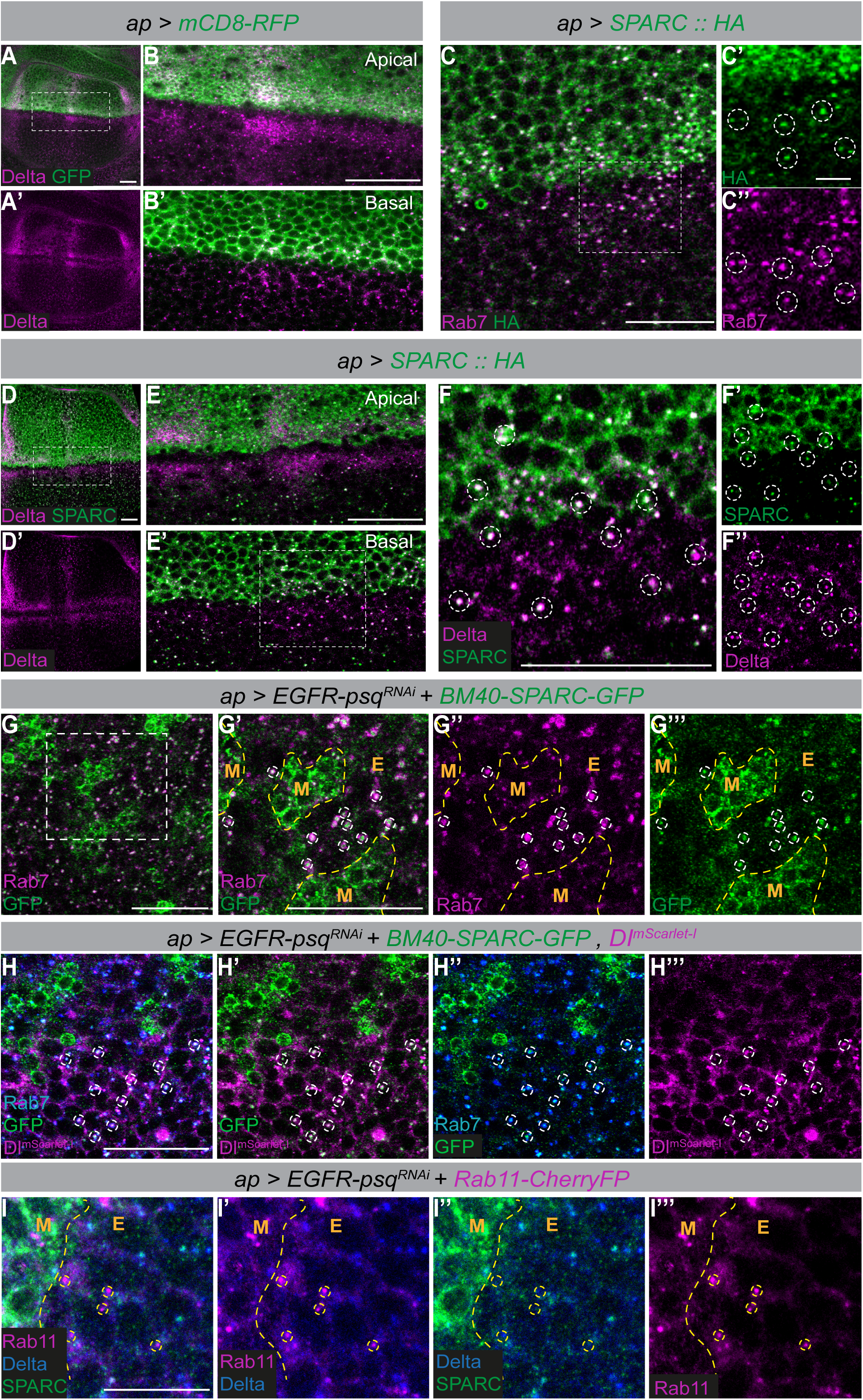
SPARC accumulates in Rab7 positive endosomes containing Delta. **(A-B’).** Expression domain of the *ap-Gal4* driver in wild-type wing discs. *ap-Gal4* (*ap>mCD8-RFP*, green) defines a restricted epithelial territory adjacent to the dorsoventral boundary, where Delta is expressed (magenta). Apical (**B**) and basal (**B’**) sections are shown. Scale bars: 25μm. **(C).** Upon epithelial overexpression of SPARC::HA (*ap>SPARC::HA*, green), SPARC localizes to Rab7-positive late endosomes (magenta). Scale bar: 25μm. Insets highlight SPARC/Rab7 positive endosomes (dashed circles). Scale bar: 10μm. **(D-F).** Upon epithelial overexpression of SPARC::HA (*ap>SPARC::HA*, green), SPARC (green, HA) accumulates in Delta endosomes (magenta). **(F-F’).** Single channels show colocalization in vesicular structures (dashed circles). Scale bar: 25μm. **(G).** *ap > EGFR-psq^RNAi^* tumors expressing BM40-SPARC-GFP (Green), Rab7 (magenta) marks late endosomes. **(G’-G’’’).** Higher magnification of boxed region in (G) shows SPARC association with Rab7 endosomes (dashed circles). Scale bar: 25μm. **(H).** *ap > EGFR-psq^RNAi^*tumors expressing BM40-SPARC-GFP (green), Rab7 (magenta), and Delta (bleu). SPARC/Rab7 endosomes are positive for Delta. Scale bar: 25μm. **(I-I’’’).** *ap > EGFR-psq^RNAi^* tumors expressing Rab11-CherryFP (magenta), Delta (bleu), and SPARC (Green). None of the Rab11 vesicles (dashed circles) colocalize with SPARC. Scale bar: 10μm.

### SPARC promotes enlargement and altered dynamics of Delta containing endosomes

The presence of SPARC within Rab7 positive endosomes raises the possibility that SPARC may alter endosomal trafficking. This also suggests that Notch signalling may be indirectly affected through altered Delta trafficking, as Delta relies on endocytic recycling for its activity [21, 22]. Furthermore, Delta is highly expressed in *EGFR-psq^RNAi^* epithelial tumour cells and is required for tumour development (Figure S2 and [23]). We observed a heterogeneous population of epithelial vesicles with variable sizes and differing levels of BM40-SPARC-GFP expression intensity in the *EGFR-psq^RNAi^* tumours (Figure 6A–A’’). Importantly, these vesicular structures, identified based on mCD8-RFP signal, correspond to endosomes, as some of them also label positive for Rab7 (Figure S2H-H’’). We therefore used the mCD8-RFP signal to segment vesicles and assess whether SPARC accumulation correlates with their size. We found a significant positive correlation between vesicle size and BM40-SPARC-GFP intensity (Spearman r = 0.61, *p < 0.0001*), indicating that larger endosomes accumulate higher levels of SPARC (Figure 6B). In addition, Rab7 positive endosomes were larger in *EGFR-psq^RNAi^*tumours compared to control wing discs (Figure 6C, E, and F). To investigate whether there is a causal link between SPARC accumulation in intracellular endosomes and their size, we overexpressed SPARC in the epithelial compartment of the wing disc, where SPARC is normally absent. This manipulation induced a significant increase in the size of Rab7 positive endosomes in the region where SPARC is overexpressed (Figure 6C–D and F), indicating that SPARC is sufficient to promote endosomal enlargement.

**Figure 6.**
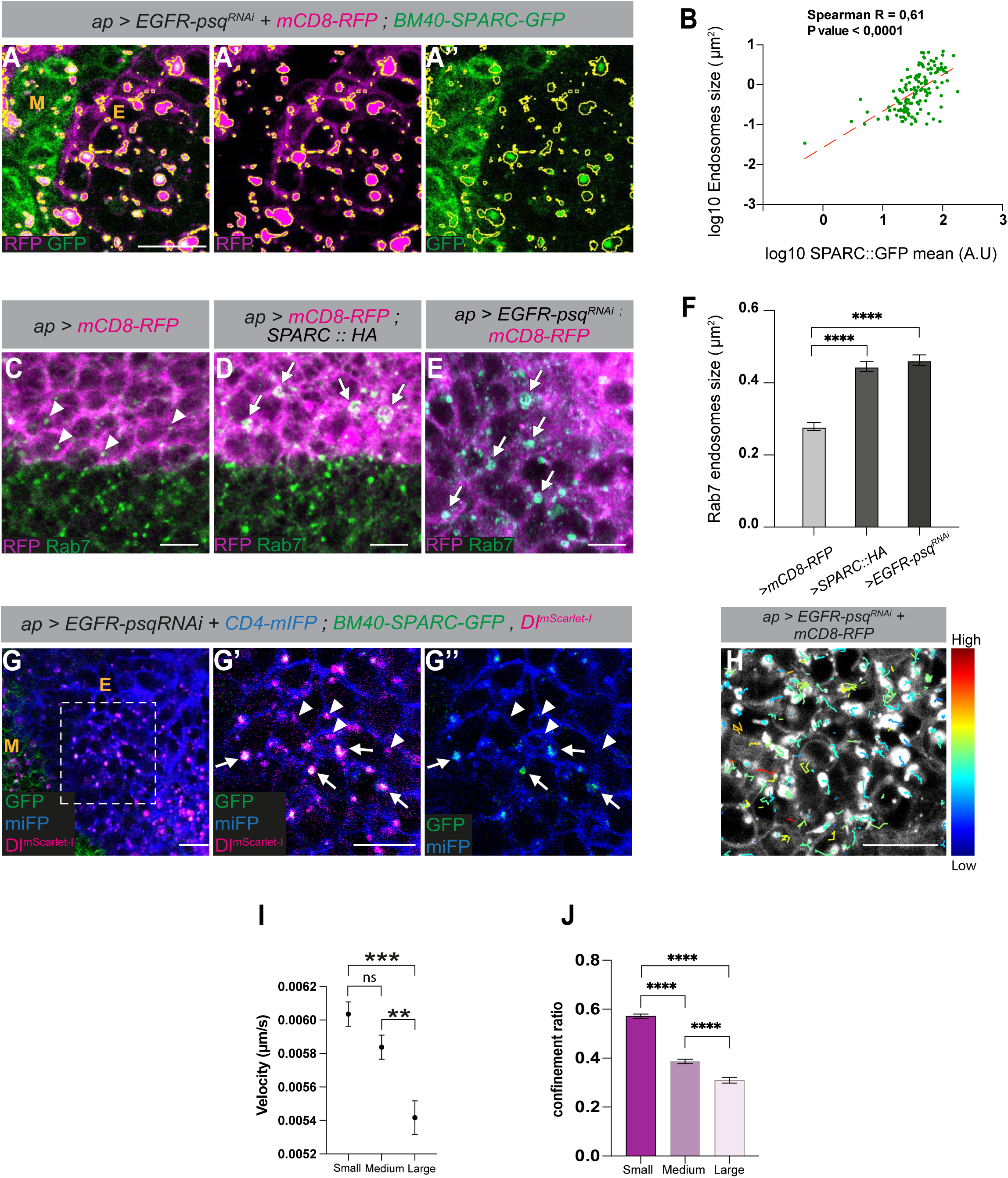
SPARC promotes endosomes enlargement and alters their motility. **(A-A’’).** *ap > EGFR-psq^RNAi^* tumours expressing *BM40-SPARC-GFP* (green) and mCD8-RFP (magenta). Epithelial vesicles are segmented according to mCD8-RFP intensity and outlined in yellow. **(B).** Scatter plot showing a positive correlation between BM40-SPARC-GFP intensity and vesicle size in *EGFR-psq^RNAi^* tumours (Spearman r = 0.61, *p < 0.0001*). Each dot represents an individual vesicle (n = 141). Data are displayed on a log10–log10 scale. A linear regression line is shown in red. **(C-E).** Rab7 endosomes (green) in control wing discs (C, ap>mCD8-RFP), upon epithelial overexpression of *SPARC::HA* (D, *ap>SPARC::HA*), and in *EGFR-psq^RNAi^* tumours (E). Epithelial cells are visualized with *mCD8-RFP* (magenta). Arrows and arrowheads show large and small endosomes, respectively. Scale bares 10μm. **(F)** Quantification of Rab7-positive endosome size in control wing discs (*ap > mCD8-RFP*), upon epithelial overexpression of *SPARC::HA* (*ap > SPARC::HA*), and in *EGFR-psq^RNAi^* tumours. Data are presented as mean ± SEM. A total of 349, 706, and 738 endosomes were analysed for control, *SPARC::HA*, and *EGFR-psq^RNAi^* tumour conditions, respectively, across three independent experiments. (Unpaired t-test, *\*\*\*\*p < 0.0001*). **(G-G’’).** Single time point from a time-lapse sequence of *ap > EGFR-psq^RNAi^* tumours expressing *BM40-SPARC-GFP* (green) and *Delta^mScarlet-I^* (magenta), together with *UAS-CD4-miFP* (blue) (see Video S4). Arrows and arrowheads show large and small endosomes, respectively. **(H).** Representative tracks of vesicles movement in tumour epithelial cells. Tracks are colour-coded according to vesicle velocity. **(I).** Quantification of vesicles velocity. (Student’s t test; \*\*\**p* < 0.001, \*\**p* < 0.01, n = 1351 small endosomes, n = 855 medium endosomes, and n = 301 large endosomes; data pooled from three independent experiments). **(J).** Quantification of endosome confinement ratio. (Student’s t test; \*\*\*\**p* < 0.0001, n = 1351 small endosomes, n = 855 medium endosomes, and n = 301 large endosomes; data pooled from three independent experiments). Scale bars: 10μm.

Given that endosome enlargement is often associated with altered endosomal properties and trafficking [24, 25], we next asked whether SPARC affects endosomal dynamics. We therefore analysed endosome motility by live imaging of *EGFR-psq^RNAi^* tumours expressing BM40-SPARC-GFP and Delta^mScarlet-I^, together with a membrane-tagged miFP to label epithelial cells (Figure 6 G-G’’). We observed heterogeneous motility behaviours among Delta-positive endosomes. Notably, enlarged endosomes were consistently SPARC-positive and exhibited reduced motility compared to smaller ones (Figure 6G–G″ and Video S4). We then tracked epithelial tumours vesicles by live imaging and compared their motility according to their size (Figure 6H and Material and Methods). Larger vesicles displayed significantly reduced velocity compared to small and medium-sized vesicles (Figure 6I). Consistently, larger vesicles exhibited a lower confinement ratio, indicating more restricted movement (Figure 6J). Together, these data indicate that SPARC accumulation within endosomes is associated with endosome enlargement and reduced motility. Such alterations in endosomal trafficking provide a mechanistic explanation to the attenuation of Notch signalling in epithelial tumour cells by mesenchymal SPARC.

### The acidic domain of SPARC is required for its localization to endosomes

As shown above, targeting of SPARC to endosomal compartments is a key step by which it modulates tumour–stroma communication. We therefore sought to identify the SPARC domain(s) responsible for its localization to these compartments. SPARC is composed of three main domains: an N-terminal acidic domain (Domain I), a follistatin-like domain (Domain II), and a C-terminal extracellular Ca²⁺-binding domain (Domain III) (Figure 7A). We used a series of HA-tagged SPARC deletion constructs lacking either individual domains or combinations of two domains [26] and expressed them in the wing disc epithelium using Ap-Gal4 driver (Figure 7A). Expression of either SPARC^DIDII^ or SPARC^DI^ resulted in colocalization with Delta positive endosomes within epithelial cells (Figure 7 B–C’’), consistent with our previous observations of the full length of SPARC (Figure 5). In contrast, constructs lacking the acidic domain (SPARC^DIIDIII^ or SPARC^DIII^) failed to accumulate in Delta-containing endosomes (Figure 7 D-E’’). These results indicate that the acidic domain of SPARC (Domain I) is specifically required for its targeting to Delta containing endosomes.

**Figure 7.**
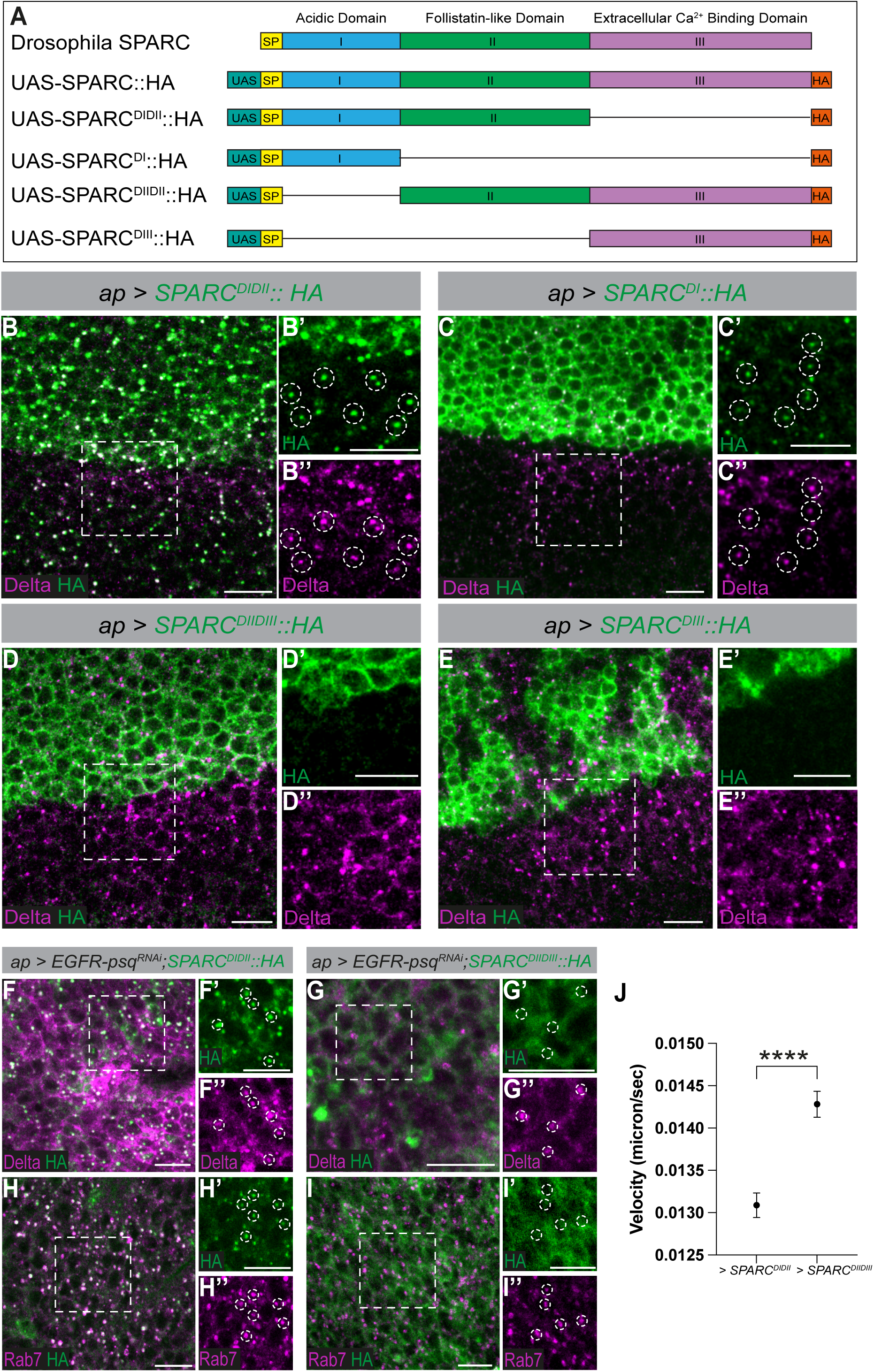
The acidic domain of SPARC is required for its targeting to Delta endosomes. **(A).** Schematic representation of SPARC domain arrangement and the HA-tagged deletion constructs. Constructs are driven by the UAS promoter (teal) and include an N-terminal signal peptide (SP; yellow), followed by Domain I (acidic; blue), Domain II (follistatin-like; green), and Domain III (extracellular Ca²⁺-binding; purple), with a C-terminal HA tag (orange). **(B-E’).** Overexpression of the *SPARC::HA* depletion constructs (green, HA) with *ap-Gal4*. Delta is indicated in magenta. Insets are higher magnification of the boxed region highlight SPARC/Delta colocalization (dashed circles). **(F-I’’).** *ap > EGFR-psq^RNAi^* tumours expressing either *SPARC^DIDII^::HA* (F-H’) or *SPARC^DIIDIII^::HA* (G-I’). Tissues were stained for HA (green), Rab7 or Delta (magenta). Insets show higher-magnification views of the boxed regions, highlighting SPARC colocalization with Delta or Rab7 endosomes (dashed circles). **(J).** Quantification of endosome velocity in tumour epithelial cells expressing SPARC deletion constructs. Endosomes in cells expressing *SPARC^DIDII^::HA* show significantly reduced motility compared to those expressing *SPARC^DIIDIII^::HA*. (****p < 0.0001, Student’s t test; n = 1474 and 1586 endosomes for *SPARC^DIDII^::HA* and *SPARC^DIIDIII^::HA* conditions, respectively; data pooled from three independent experiments). Scale bars: 10 μm.

We next investigated whether the requirement of the acidic domain of SPARC for endosomal entry is maintained in the *EGFR-psq^RNAi^* tumor context. To do so, we expressed two SPARC deletion constructs, SPARC^DIDII^, which retains the acidic domain, and SPARC^DIIDIII^, which lacks it, in epithelial tumor cells of *EGFR-psq^RNAi^* discs. Consistent with our previous observations, SPARC^DIDII^ robustly colocalized with both Delta positive and Rab7 positive endosomes within epithelial tumor cells (Figure 7 F–H’’). In contrast, SPARC^DIIDIII^ failed to accumulate in Delta or Rab7 positive endosomes and instead displayed a diffuse distribution (Figure 7G–I’’), confirming that the acidic domain is required for SPARC targeting to endosomes in the tumor epithelium. Furthermore, live imaging of the endosome’s motility revealed that endosomes in epithelial tumor cells expressing SPARC^DIDII^ exhibited significantly reduced velocity compared to endosomes in cells expressing SPARC^DIIDIII^ (Figure 7J). Together, these data showed that the acidic domain of SPARC is required both for its localization to Rab7 positive endosomes and for its ability to modulate endosomal trafficking features in epithelial tumor cells.

## Discussion

In this study, we describe a mechanism by which SPARC functions as an intercellular mediator of tumour–host communication. We show that SPARC, produced by mesenchymal cells, is internalized by the *EGFR-psq^RNAi^* epithelial tumour cells and trafficked to Rab7-positive late endosomal compartments, where it colocalizes with the Notch ligand Delta. In *EGFR-psq^RNAi^* tumours, SPARC alters endosomes properties resulting in enlargement and reducing motility. These alterations indirectly reduce Notch signalling (Figure 8).

**Figure 8.**
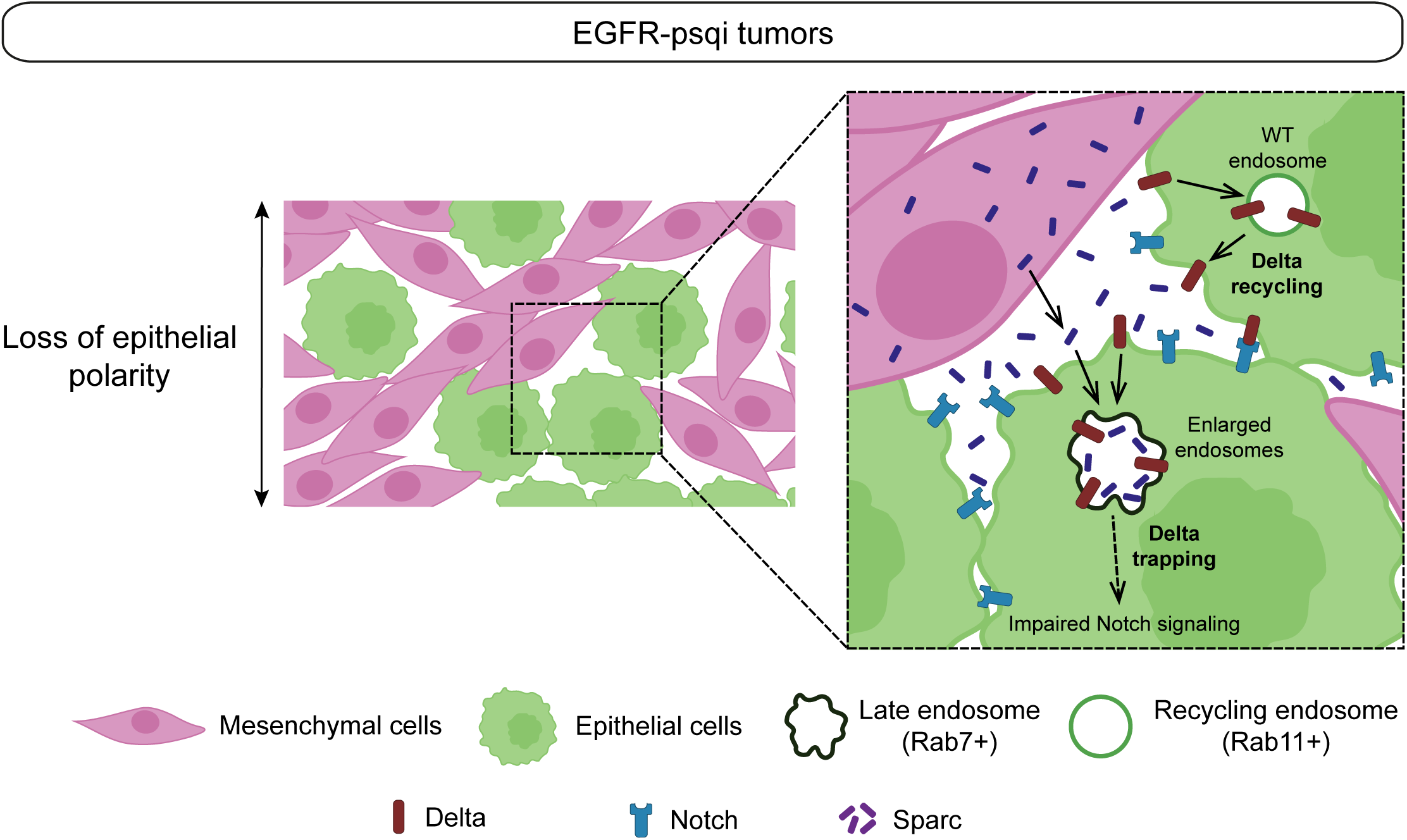
Model for SPARC mediated regulation of Delta trafficking and Notch signalling in *EGFR-psq^RNAi^*tumours. Mesenchymal cells produce and secrete SPARC, which is transferred to neighboring epithelial tumour cells. SPARC accumulates in Rab7 positive late endosomes, promoting endosomal enlargement and trapping of the Notch ligand Delta. This prevents efficient Delta recycling through Rab11-positive compartments and results in reduced Notch signalling output in tumour epithelial cells.

The localization of SPARC within endosomal compartments is unexpected for a matricellular protein. Interestingly, studies of human ovarian cancer have shown that SPARC protein accumulates in the cytoplasm of malignant epithelial cells despite its mRNA being detected exclusively in the surrounding stromal cells [27]. In contrast, SPARC is absent from epithelial cells in benign ovarian tumours, where an intact basement membrane separates the stromal and epithelial compartments. These observations led to the proposal that the loss of the basement membrane in malignant tumours permits stromal-derived SPARC to access and become internalized by neighbouring cancer cells [27]. An interesting parallel can be drawn with the *EGFR-psq^RNAi^* wing disc tumour model. In wild-type wing discs, SPARC is produced by the associated myoblasts but is not detected within the epithelial compartment, consistent with the separation of these tissues by an intact extracellular matrix. In contrast, *EGFR-psq^RNAi^* tumours disrupt tissue architecture, leading to intermingling of epithelial and mesenchymal cells and extensive extracellular matrix remodelling. Under these conditions, SPARC is efficiently transferred from myoblasts to epithelial tumour cells, where it accumulates in endosomal compartments.

Tumour growth often depends on coordinated interactions between cancer cells and surrounding stromal cells, which co-evolve during progression and contribute to tumour expansion and behaviour [3, 28]. This synergy is well illustrated in the *EGFR-psq^RNAi^* model [12, 13]. In this model, epithelial tumour cells and tumour-associated mesenchymal cells exchange intercellular signals and expand to comparable sizes despite originating from tissues with distinct growth properties. In the wild-type wing disc, epithelial cells are substantially larger than the associated myoblasts, whereas in *EGFR-psq^RNAi^* tumours the two populations become similar in size [12, 13]. This shift suggests that mechanisms operate to coordinate the expansion of these two compartments during tumour development. Our findings raise the possibility that mesenchymal SPARC contributes to this coordination, at least in part by attenuating epithelial Notch signalling, thereby restraining epithelial growth and allowing the mesenchymal compartment to expand in parallel.

Alterations in endosomal trafficking influence signalling activity and intercellular communication during tumour growth. For example, in *Drosophila*, mutation of *vps25* - a key component of the ESCRT endosomal sorting machinery - leads to the accumulation of the Notch receptor within enlarged endosomal compartments [24, 29]. This results in constitutive Notch signalling, ectopic expression of the secreted ligand Unpaired (Upd), and subsequent activation of the JAK–STAT pathway in neighbouring wild-type cells, driving non-autonomous tissue over proliferation. Our findings reveal a distinct mechanism by which endosomal trafficking modulates tumour development. We find that SPARC accumulates in Rab7-positive late endosomes alongside the Notch ligand Delta. This accumulation correlates with reduced Notch activity in epithelial tumour cells. These contrasting outcomes underscore the context-dependent nature of endosomal regulation: while trapping the Notch receptor in endosomes promotes ligand-independent activation, promoting the ligand Delta within late endosomes may limit its recycling to the plasma membrane, and consequently attenuating Notch signalling.

SPARC has been widely implicated in tumour–stroma interactions and is frequently expressed by stromal or immune cells, where it influences tumour progression, invasiveness, and patient outcome across multiple cancer types [9, 30, 31]. Despite this clinical relevance, the molecular mechanisms by which stromal-derived SPARC signals to tumour cells and modulates intracellular signalling pathways have remained elusive. Moreover, SPARC has been reported to exert context-dependent functions, acting either as a tumour promoter or tumour suppressor depending on tissue type and tumour stage [8]. This functional duality has complicated efforts to define the mechanistic basis of SPARC activity. Our findings reveal that SPARC can act as an intracellular regulator of signalling by modulating endosomal trafficking. Specifically, SPARC accumulates in Rab7-positive late endosomes and shows no detectable association with Rab11-positive recycling endosomes. Its accumulation in this compartment is associated with its enlargement, a phenotype linked to defective endosomal trafficking. Delta activity depends on its trafficking between endosomal compartments and the plasma membrane to sustain Notch signalling. We propose that, when localized to SPARC-positive endosomes, Delta is preferentially directed toward late endosomal compartments rather than recycled back to the plasma membrane. Such misdirection would limit Delta availability for signalling and thereby attenuate Notch pathway activation. Given the central role of Notch signalling in cancer [32], this modulation of Delta intracellular trafficking constitutes a key regulatory point and may represent a broadly relevant form of tumour–stroma communication and may help explain the context-dependent effects of mammalian SPARC–Notch interactions at tumour–host interfaces.

## Materials and Methods

### Drosophila melanogater strains and genetics

All *Drosophila melanogaster* stocks were grown on standard medium at 25°C. The following stains were used: *w^1118^* as wild type (wt), UAS-mCD8GFP (BL#5137), UAS-mCD8-RFP (BL# 27391), UAS-mCD8-RFP (BL# 27392), Ap-Gal4 (BL#25685), SPARC^[MI00329-GAL4]^ (BL#77473), BM40-SPARC-GFP [17], 15B03-LexA [12], lexAop-CD8-GFP (BL# 66545), *EGFR-psq^RNAi^* [12], UAS-SPARC::HA, UAS-SPARC^DI^::HA, UAS-SPARC^DIDII^::HA, UAS-SPARC^DIIDIII^::HA, UAS-SPARC^DIII^::HA [26], Notch [ts1] (BL# 2533), NRE-RFP [33], Ubiquitin-Rab11cherryFP [34], Delta^mScarlet-I^ [13], UAS-CD4-miFP (BL# 64183), CG9650-GFP (CPTI#1741), Df(3R)BSC524 (BL#25052). The *EGFR-psq^RNAi^* tumours were inducted as described in [12].

### Generation of the *LexAop-sparc^RNAi^* fly line

Inverted repeat fragments targeting *sparc*, corresponding to the sequence used in the *UAS-sparc^RNAi^* line (VDRC kk108185), were cloned into the pJFRC19 plasmid [35] using the NotI and XbaI restriction sites. The resulting *LexAop-sparc^RNAi^* construct was integrated into the attP landing site at 68A4 on chromosome III via φC31-mediated transgenesis following embryo injection into *nos-phiC31-NLS; attP2* flies. Knockdown efficiency was validated by expressing *LexAop*-*sparc^RNAi^* in mesenchymal cells using *15B03-LexA* and assessing SPARC protein in *EGFR-psq^RNAi^* tumours (Figure S2F-G’).

### Immunostainings and in situ hybridization

Preparation of the *EGFR-psq^RNAi^* and immunofluorescence of the wing discs was performed as described in [12, 18]. The following primary antibodies were used for immunofluorescence staining: Chicken anti-GFP (1:300, Abcam, ab13970), Rabbit anti-SPARC (1:500, Maurice Ringuette, University of Toronto, Canada), Rat anti-RFP (1:800, Chromotek, 5F8-20), Mouse anti-Cut (1:10,DHSB, 2B10), Rat anti-DCAD2 (1:200, DHSB, DCAD2-s), Mouse anti-Delta (1:50, DHSB, C594.9B), Mouse anti-NICD (1:100, DSHB), Mouse anti-Rab7 (1:25, DHSB, Rab7-s) Anti-HA (1:1000, Roche, clone 3F10), Rabbit anti-Twist (1:2000, Eileen Furlong, Heidelberg, Germany). Single-molecule fluorescence in situ hybridization (smFISH) experiments were performed as described in [18]. *Delta* and *sparc* probe sets were generated by Biosearch Technologies. Probe sequences targeted exon 3 of each gene.

### Genetic rescue of *sparc* loss-of-function lethality

To assess the functionality of the BM40-SPARC-GFP cassette, a genetic rescue experiment was performed using the *sparc* loss-of-function allele *SPARC ^[MI00329-GAL4]^*. Flies carrying the BM40-SPARC-GFP transgene were first recombined with SPARC^[MI00329-GAL4]^ to generate the stock BM40-SPARC-GFP, SPARC^[MI00329-GAL4]^. Virgin females from this stock were crossed to males carrying the deficiency Df(3R)BSC524/TM6C, Sb, which deletes a couple of genes including *sparc*. Progeny were scored for the presence of the BM40-SPARC-GFP, SPARC^[MI00329-GAL4]^/Df(3R)BSC524 genotype. In the absence of rescue, this genotype is expected to be lethal due to loss of *sparc*. Recovery of viable adults with this genotype indicates functional complementation by the BM40-SPARC-GFP cassette. Genotype frequencies among the progeny were compared with expected Mendelian ratios to evaluate rescue efficiency (Figure S3).

### Quantitative RT PCR

For each condition, 30–40 wing discs were dissected and total RNA was extracted using TRIzol reagent (Life Technologies), according to the manufacturer’s instructions. RNA was reverse-transcribed into cDNA using M-MLV reverse transcriptase (Promega, M531A). Quantitative PCR was performed on a LightCycler 480 system using SYBR Green I Master Mix (Roche, 04707516001). Three independent biological replicates were analysed for each condition. Transcript levels were normalized to the housekeeping gene Rpl32.

### Microscopy and data analysis

#### Fixed samples

Fluorescent samples were imaged at the BIOSIT facility (University of Rennes) using either an LSM980 Airyscan2 or a Leica SP8 confocal microscope. Images were acquired with 20×, 40×, or 63× objectives, at a frame size of 1024 × 1024 pixels. Image processing was performed using ImageJ, and final figures were assembled in Adobe Illustrator. For fluorescence quantifications, all experimental conditions were processed and analyzed in parallel. Confocal images were acquired using identical laser power and detector settings across samples. For each z-stack, sum-intensity projections were generated prior to measurement.

#### Live imaging

Larvae were dissected in Schneider’s Drosophila medium. Tumour-bearing tissues were mounted in Schneider’s medium supplemented with penicillin–streptomycin (Invitrogen 15140-122), 15% fetal bovine serum (Biosera FB-1200/500), and insulin (Sigma I5500). Samples were mounted between an 18 × 18 mm coverslip (VWR 631-1567) and a 35 mm Lumox dish (Sarstedt 94.60077.331). To prevent evaporation during acquisition, preparations were covered with Halocarbon oil 27 (Sigma H8773). Live imaging was performed on an LSM980 Airyscan2 microscope (BIOSIT, University of Rennes) with the chamber maintained at 25°C. Time-lapse sequences were acquired using a 63× objective, at a resolution of 1024 × 1024 pixels, with images collected every 30 s.

For endosomal BM40-SPARC-GFP intensity quantification (Figure 6A–B), wing discs expressing *EGFR-psq^RNAi^*, BM40-SPARC-GFP, and the membrane marker mCD8–RFP were prepared and imaged by confocal microscopy. Endosomal structures were segmented automatically in Fiji/ImageJ using the thresholding detector implemented in the TrackMate plugin, based on the mCD8–RFP signal. The mean of BM40<u>-</u>SPARC<u>-</u>GFP fluorescence intensity was subsequently measured within the corresponding regions. Correlation between endosome area and BM40<u>-</u>SPARC<u>-</u>GFP levels was assessed using Spearman’s rank correlation coefficient, and the regression trend is shown in the scatter plot.

For endosome tracking motility analysis, endosomes were detected and tracked over time using the TrackMate plugin in Fiji/ImageJ, generating trajectories assigned to unique Track IDs. TrackMate derived motility parameters, including mean speed, confinement ratio, were extracted and matched to the corresponding Track IDs. Endosomes were then classified into three size groups based on mean area (small < 0.2 µm², medium 0.2–0.5 µm², and large > 0.5 µm²). Distributions of motility parameters were compared between size classes using non-parametric statistical tests and visualized with plots.

## Acknowledgments

We would like to thank the Bloomington Drosophila Stock Center, the Vienna Drosophila Resource Center (VDRC), and the Developmental Studies Hybridoma Bank for providing *Drosophila* strains and antibodies. We are grateful to Roland Le Borgne, Peter Lorincz and Caroline Dillard for the critical reading of the manuscript. We acknowledge Xavier Pinson and Stephanie Dutertre from the Microscopy Rennes Imaging Center (MRic, BIOSIT, Biogenouest) for technical assistance, and Cyrille Surbled for fly stock maintenance and technical help. Work in the Bray lab was supported by a program grant from the Medical Research Council (MR/L007177/1 and MR/T014156/1). Work in the Boukhatmi lab was supported by an AFM-Téléthon Trampoline Grant (#23108), ATIP-Avenir (CNRS), and la Ligue contre le Cancer. N.A. was supported by an AFM-Téléthon PhD fellowship (#23846) and an FRM fellowship (FDT202404018608). E.L was supported by an AFM-Téléthon PhD fellowship (#29992). T.M. is supported by the Fundação para a Ciência e a Tecnologia (FCT) CEEC 5th Edition Assistant Researcher Award (DOI: 10.54499/2022.02406.CEECIND/CP1720/CT0014).

---

**Figure S1.**
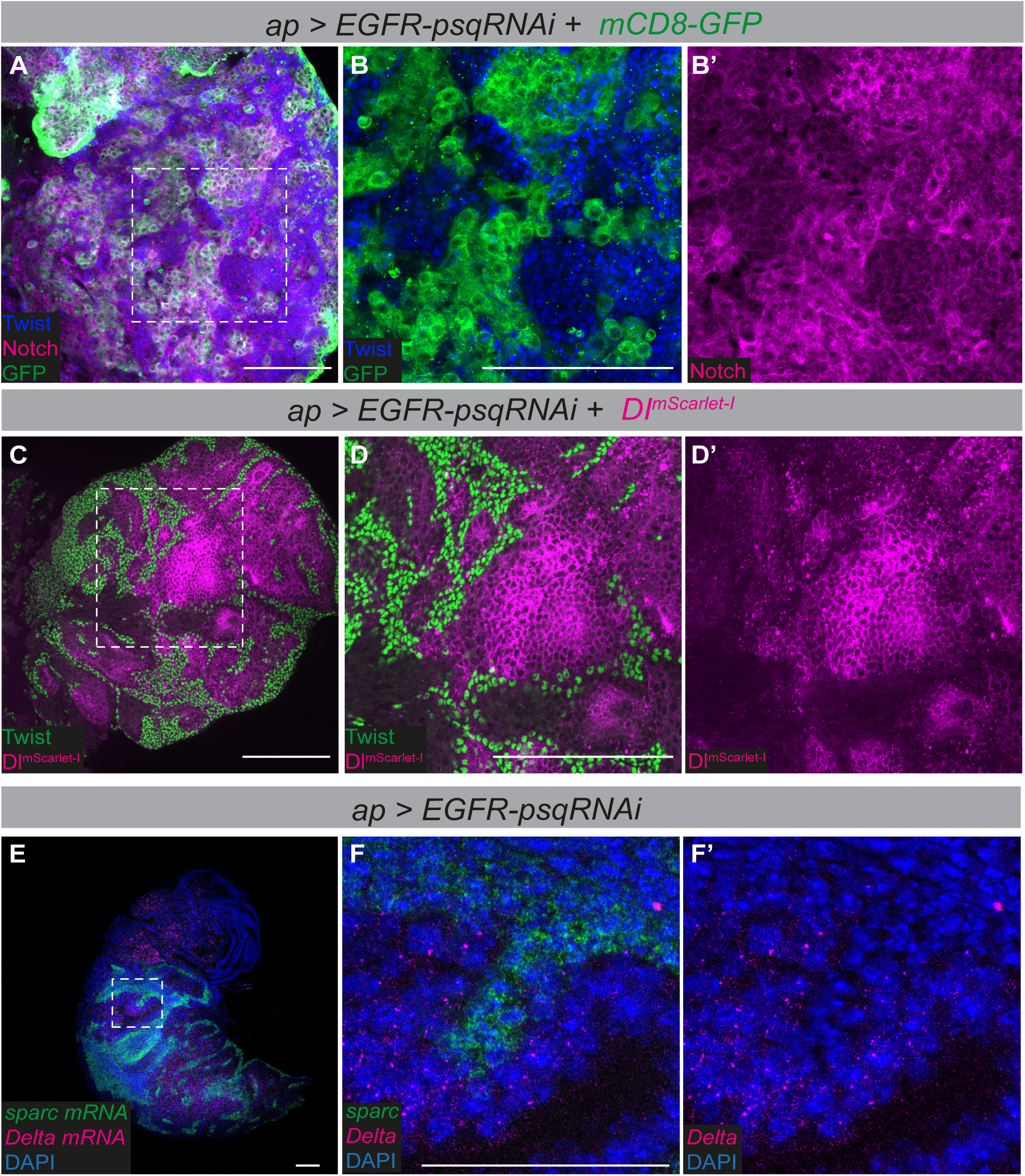
Distribution of the receptor Notch and the ligand Delta in the *EGFR-psq^RNAi^* tumour. **(A-B’).** Tumour induced by the expression of EGFR and *psq^RNAi^* (*ap-Gal4 > UAS EGFR-psq^RNAi^*) co-expressing mCD8-GFP. **(B-B’).** Higher magnifications of the boxed region in A. GFP (green) marks the epithelial cells, Twist (blue) the mesenchymal cells. Notch (magenta) is expressed in both the epithelial and the mesenchymal cells. Scale bars: 100 μm. **(C-D’).** *EGFR-psq^RNAi^* tumours co-expressing Dl^mScarlet-I^. Dl^mScarlet-I^ ligand (magenta) is homogeneously detected in the epithelial cells. The mesenchymal cells are marked by Twist (green). **(D-D’).** Higher magnification of the boxed region in (C). Scale bars: 100 μm. **(E-F’).** *Delta* mRNA transcripts (magenta) are detected in the epithelial cells. *sparc* mRNA (green) are detected in the mesenchymal cells. DAPI (blue) marks nuclei. **(F-F’).** Higher magnification of the boxed region in F. Scale bar: 50 μm.

**Figure S2.**
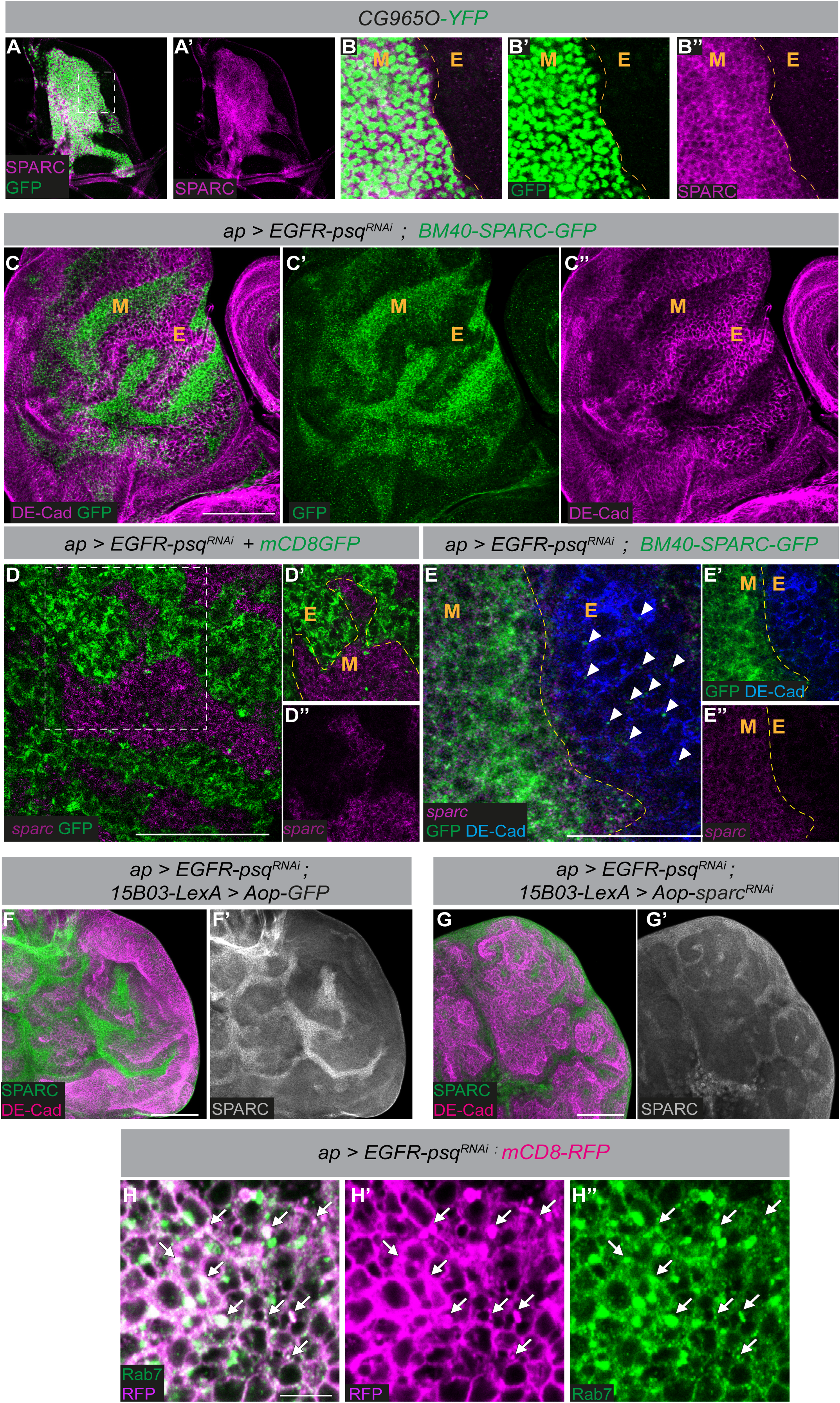
Expression of SPARC in wild type wing discs and *EGFR-psq^RNAi^* tumours. **(A-A’).** SPARC (magenta) is detected in the wing disc associated myoblasts (green, *CG965O-YFP*). **(B-B’’).** Higher magnifications of the boxed region in A. Scale bars: 50 μm. **(C-C’’).** BM40-SPARC-GFP (green) is detected in both the mesenchymal (M) and the epithelial cells (E) of the *EGFR-psq^RNAi^* tumours. DE-Cad (magenta) marks the epithelial cells. Scale bar: 100 μm. **(D).** *sparc* mRNA transcripts (magenta) are detected in mesenchymal cells (M) and are absent from epithelial cells (E) of *EGFR-psq^RNAi^*tumours, which are visualized by mCD8-GFP (green; *ap-Gal4, UAS-mCD8-GFP*). **(D’-D’’).** Cropped views of the boxed region in (D) showing *sparc* transcript distribution. Scale bar: 50 μm. **(E-E’’).** Analysis of *sparc* mRNA transcripts (magenta) and BM40-SPARC-GFP (green) in *EGFR-psq^RNAi^* tumour. BM40-SPARC-GFP (green) is detected in both mesenchymal cells (M) and epithelial cells (E), marked by DE-Cad (blue, arrows) (E). Yellow dashed lines delineate mesenchymal (M) and epithelial (E) compartments. Scale bar: 50 μm. **(F-G′).** Validation of *sparc* knockdown efficiency in *EGFR-psq^RNAi^* tumours. Control tumours (*15B03-LexA; Aop-GFP*; F–F′) show SPARC signal in the mesenchymal cells. Tumours expressing *sparc* RNAi in mesenchymal cells (*15B03-LexA; Aop-sparc^RNAi^*; G–G′) show a reduction of SPARC levels. SPARC is shown in green and grayscale. DE-Cad (magenta) marks epithelial tumour cells. Scale bar: 100 μm. **(H–H″).** in *EGFR-psq^RNAi^*tumours (ap > EGFR-psqRNAi; mCD8-RFP) shows that vesicular structures identified by mCD8-RFP (magenta) colocalize with the late endosomal marker Rab7 (green, arrows). Merge (H), RFP channel (H′), and Rab7 channel (H″) are shown. Scale bar: 5 μm.

**Figure S3.**
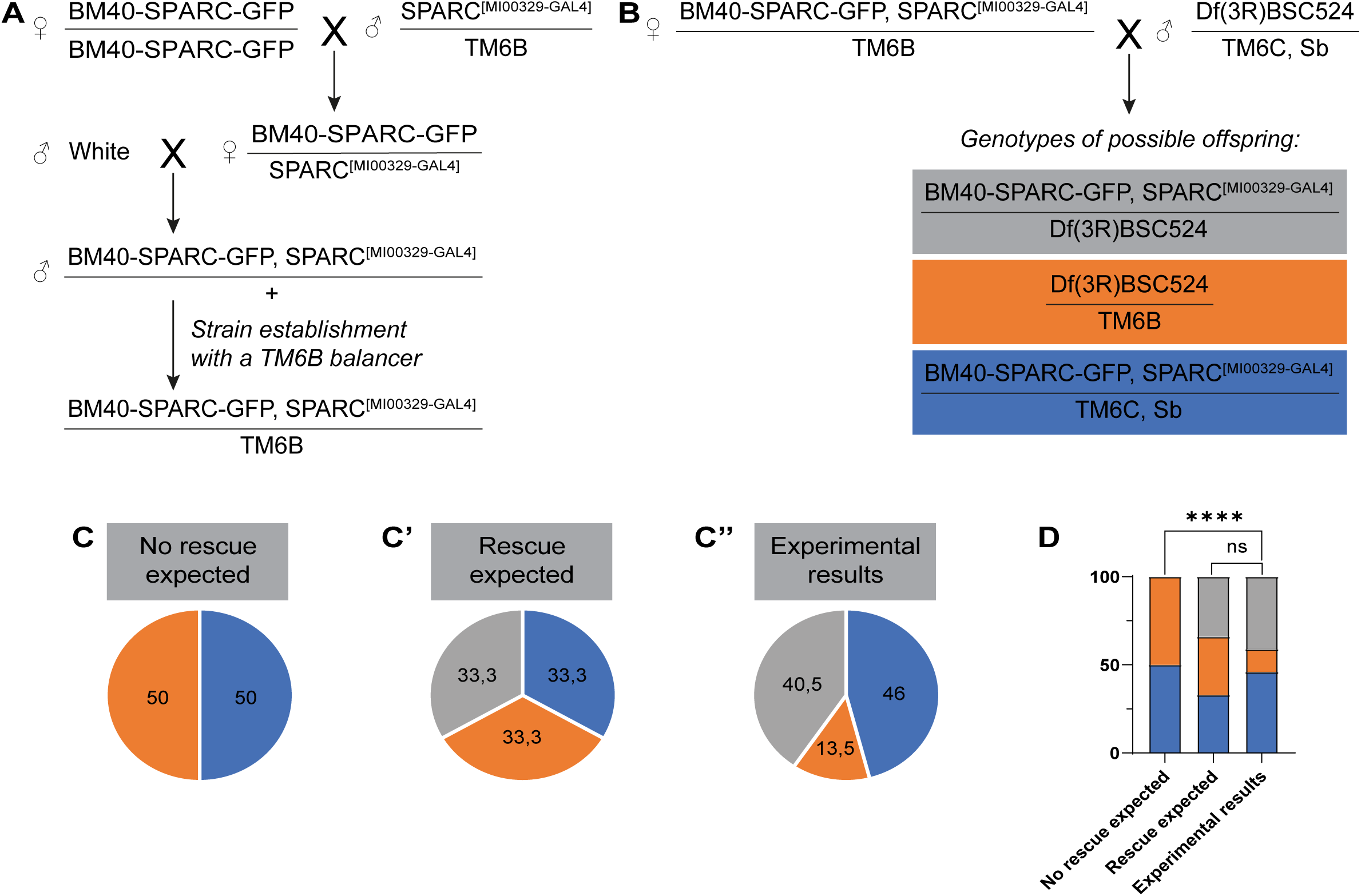
Functional validation of the BM40-SPARC-GFP cassette by genetic rescue of *sparc* loss-of-function lethality. **(A).** Genetic scheme used to generate flies carrying both the *BM40-SPARC-GFP* cassette and the *sparc* loss-of-function allele (*SPARC ^[MI00329-GAL4]^*) balanced over TM6B. **(B).** Cross performed to test rescue of the *sparc* mutant phenotype using the deficiency Df(3R)BSC524, which removes the sparc locus. The possible genotypes expected among the progeny are indicated. **(C-C″).** Comparison between expected Mendelian genotype distributions and experimentally observed progeny. **(C)** Expected ratios in the absence of rescue. **(C′)** Expected ratios if the BM40-SPARC-GFP cassette fully rescues the mutation**. (C″)** Experimentally observed genotype frequencies obtained from the cross shown in (B). **(D).** Quantification of genotype distributions comparing theoretical expectations with experimental results. Coloured bars indicate the proportion of each genotype. The observed distribution closely matches the predicted rescue scenario, indicating that the BM40-SPARC-GFP cassette restores viability of *sparc* mutant animals. (\*\*\*\**p* < 0.0001, Chi2 Test)

